# Protection of phage applications in crop production: a patent landscape

**DOI:** 10.1101/555904

**Authors:** D. Holtappels, R. Lavigne, I. Huys, J. Wagemans

## Abstract

In agriculture, the prevention and treatment of bacterial infections represents an increasing challenge. Traditional (chemical) methods have been restricted to ensure public health and limit the occurrence of resistant strains. Bacteriophages could be a sustainable alternative. A major hurdle towards the commercial implementation of phage-based biocontrol strategies concerns aspects of regulation and intellectual property protection. Within this study, two datasets have been composed to analyze both scientific publications and patent documents and to get an idea on the focus of R&D by means of an abstract and claim analysis. 137 papers and 49 patent families were found searching public databases, their numbers increasing over time. Within this dataset, the majority of the patent documents were filed by non-profit organizations in Asia. There seems to be a good correlation between the papers and patent documents in terms of targeted bacterial genera. Furthermore, granted patents seem to claim rather broad and cover methods of treatment. This review shows that there is indeed a growing publishing and patenting activity concerning phage biocontrol. Targeted research is needed to further stimulate the exploration of phages within integrated pest management strategies and to deal with bacterial infections in crop production.

## 1. Introduction

Soon after the discovery of bacteriophages by d’Herelle and Twort at the beginning of the 20^th^ century [1,2], the potential to use these bacterial viruses as a therapeutic was recognized. Although the first applications of phages focused on human medicine [3], other fields including agriculture soon began to explore the potential of bacteriophages as biocontrol agents [4]. The first isolation of phages infecting plant pathogenic bacteria (PPB) dates back to 1924, when it was shown that *Xanthomonas campestris* pv. *campestris*, causing black rot in *Brassicaceae,* could be lysed by the filtrate of diseased cabbages [4,5]. The following years, interest in phages as biocontrol agents remained relatively high [4]. However, the discovery of broad-spectrum antibiotics and other bactericidal chemicals resulted in a dwindling interest in phage therapy research in general [6].

Within the agricultural sector, the prevention and treatment of bacterial infections represents an increasing challenge. For farmers, devastating losses by bacterial pathogens are generally estimated to reach 10% of the total production [7] and for some bacterial species like *Xanthomonas campestris* pv. *campestris* crop yield can be reduced by 25% [8]. Other major threats include *Ralstonia solanacearum, Xylella fastidiosa* and *Pseudomonas syringae* pathovars [9]. In recent years, general antibiotics like streptomycin as well as copper-based chemicals have been restricted in crop production to ensure public health and limit the occurrence of resistant strains [10–13]. Because of these restrictions, the search for sustainable, natural biocontrol of PPBs has come to a critical stage, especially considering the increased food production needs [14]. Governments have decided to implement Integrated Pest Management strategies (IPMs) as the standard for crop protection (e.g. The European Parliament and the Council of the European Union (2009) Directive 2009/128/EC of the European Parliament and of the Council of 21 October 2009) [15]. These strategies are based on the implementation of sustainable pest control strategies with the emphasis on biological control not to eradicate pests, but to maintain their populations to avoid economical losses [15]. As phages are the natural predators of bacteria, and thus fit into this framework of IPM, research on phage-based biocontrol is getting back into the picture. Some critical proofs of concept on the efficacy of phages as biocontrol agents have been demonstrated in recent years as reviewed elsewhere [4].

A major hurdle towards the commercial implementation of these phage-based biocontrol strategies in crop protection concerns aspects of regulation and intellectual property protection. The cost to develop new crop protectants can go up to $286 million and may take eleven years [16,17]. Therefore, patents can act as a tool to stimulate innovations in the field. They provide the applicant(s) the right to prohibit others to use their invention for a time span of twenty years in a specific geographical area. This protection could assure a return-on-investment to the patent holder and hence serve as an incentivizing tool for research and innovation [18,19]. However, patenting biological substances like bacteriophages, has been shown to be difficult [20,21]. Nevertheless, the question remains whether phages and phage-containing products are protectable by patents and what the scope is of these patents. Here, we present a patent landscape on the existing patents within the field of phage biocontrol using natural phages in agriculture and correlate this with a systematic survey of the scientific literature. A database containing the relevant patent documents within this research area has been set up and analyzed. The number, geographical distribution and legal status were examined and the scope of the patents was investigated by means of a claim analysis. In parallel, a database of scientific publications within the same research topic was analyzed to enable comparisons with the patent information. This analysis allows us to draw conclusions on the amount and scope of protection of phage-based biocontrol applications in crop production.

## 2. Methodology

### Dataset scientific publications

A dataset consisting of scientific publications on the topic of phage biocontrol in crop production has been created by searching Web of Science and PubMed using Boolean search operators combined with a set of keywords relevant for this topic (((“phage”[Title/Abstract] OR “phage therapy” [Title/Abstract] OR “bacteriophage” [Title/Abstract] OR “phage biocontrol” [Title/Abstract] OR “bacteriophage therapy” [Title/Abstract])) AND ((“plant”[Title/Abstract] OR “crop”[Title/Abstract])). The database has been manually curated by eliminating non-relevant papers (papers discussing human phage therapy or phage biocontrol in food). Therefore, all abstracts of the scientific publications have been manually curated to verify the relevance to the topic. The last update of the dataset was on the 17th of February 2019 (Supplementary Table S1) and can be considered as an up-to-date snapshot of the situation.

### Patent search, legal status and geographical distribution

An algorithm based on Verbeure *et al.* (2006) [22] was used to set up a dataset of relevant patent applications and granted patents. In short, a classification search was performed using the International Patent Classification system (IPC) (A01N63 – C12N7) combined with a set of keywords relevant for phage applications in the agricultural sector (bacteriophage, phage, phage biocontrol, phage therapy, bacteriophage therapy, plant, crop) in public databases (EspaceNet, Patent Scope, Google Patents). The dataset was manually evaluated and was last updated the 17th of February 2019 (Supplementary Table S2).

The patent landscape was performed according to Huys *et al.* (2009) [23]. Three different categories of the legal status were used as retrieved from EspaceNet by evaluating the “Global dossier” in case of non-European patents or the “INPADOC legal status” and “EP register” for European patents: Pending (patent application in consideration), Granted (patent that is active in a specific territory) and Dead, meaning that the application or patent is abandoned (e.g. WO2014177996 (A1)), refused (e.g. JP2005073562 (A)), withdrawn or deemed to be withdrawn (e.g. CN103430973 (A)) or lapsed (e.g. US19840662065). A patent and/or application is considered “active” when the application is still pending or when the patent is granted. On the contrary, if the patent and/or application is dead, it is considered “non-active”.

The applicants and the geographical span of the patents were evaluated by looking at the patent document and its attributed number. The documents were categorized according to continent to have a better overview (Africa (ZA), Asia (CN, IN, JP, KR), Oceania (AU, NZ), Europe (DE, EA, EP, ES, GB, HU, IT), North America (CA, US), South America (AR, BR, CL, CR, GT, MX, PE)).

### Categorization of patent documents and scientific publications

To have an understanding of the core focus of patent documents and scientific publications in terms of genera of bacteria being addressed, both have been categorized among the most prominent bacterial genera causing plant bacterial diseases as determined by Mansfield *et al.* 2012: *Agrobacterium, Dickeya, Erwinia, Pectobacterium, Pseudomonas, Ralstonia, Xanthomonas, Xylella* and Other [9]. Here, “Other” has been defined as any other bacterial genus or if the document did not specify the bacterium addressed. The categorization was based on a claim analysis (both dependent and independent claims) of the patent documents and an abstract analysis of the scientific publications.

A detailed independent claim analysis has been performed for the granted, active patents to determine the scope of the patent. Independent claims were analyzed as these generally define the broadest scope. Claims have been categorized into four categories: (i) the phage itself (“phage”), (ii) the composition of the phage cocktail and the final product (“composition”), (iii) methods for producing and/or obtaining phages (“production”) and (iv) the use of phages to treat (non-human) bacterial diseases (“treatment”). One limitation of this research is that the authors had to rely on automatic translations of the claims in Asian patents in order to understand their scope. The same categories have been applied on the scientific publications by interpreting the abstracts of the papers, enabling a comparison between the scope of patents and scientific papers.

Moreover, the impact of the independent claims has been evaluated based on Huys *et al.* 2009 [23] according to Art. 69, Art. 83 and the Protocol of the European Patent Convention (EPC) (“fair protection for the patentee with a reasonable degree of certainty for third parties”). In case of US patents, the claim interpretation was based on US Utility Patent Act §112, demanding a “clear written description” and “best mode for carrying out the invention”. Non-European and non-US patents were interpreted by the authors in a similar way. Using this methodology, three impact levels could be defined: narrow, intermediate and broadly defined claims. The circumvention of the claims was estimated according to the author’s appraisal. Narrow defined claims (green) cover specific details of the invention and can easily be circumvented e.g. changes in the genomic sequence of the phage, adaptations to the product composition, different protocols and methods of treatment are not covered by the claims. Intermediate claims (orange) cover the invention as such without describing details (e.g. a specific phage for a specific bacterium, a composition of a product or production process). These claims can be circumvented though this would require substantial inventiveness. The authors have categorized production and treatment claims as intermediate when these claims cover broad methods of production of treatment but only for a particular phage. On the other hand, broadly defined claims (red) cover every aspect of the invention, but are vaguely described and hence claim outside the invention. However, no full freedom-to-operate analysis for each individual invention has been made as this is beyond the scope of this study.

## 3. Results

### Patent search, scientific papers and legal status

Searching the different databases resulted in the identification of 49 different patent families within the field of phage biocontrol in crop production. A patent family is defined as a group of patents and/or patent applications that have been filed in different countries but protect one and the same invention and have the same inventor(s). To highlight specific patent families, the first attributed application number has been chosen to represent the family. Within the database, the families comprise in total 97 patents and applications (both active and non-active). Figure 1 shows the different patent families organized per year (priority year – dark blue area chart). It shows that the number of patent families slightly increases in time (peak at 2013) and decreases again as less patents have been filed in 2016. Information on patent applications from 2017, 2018 and 2019 is not complete as these applications may not have been published yet.

**Figure 1.**
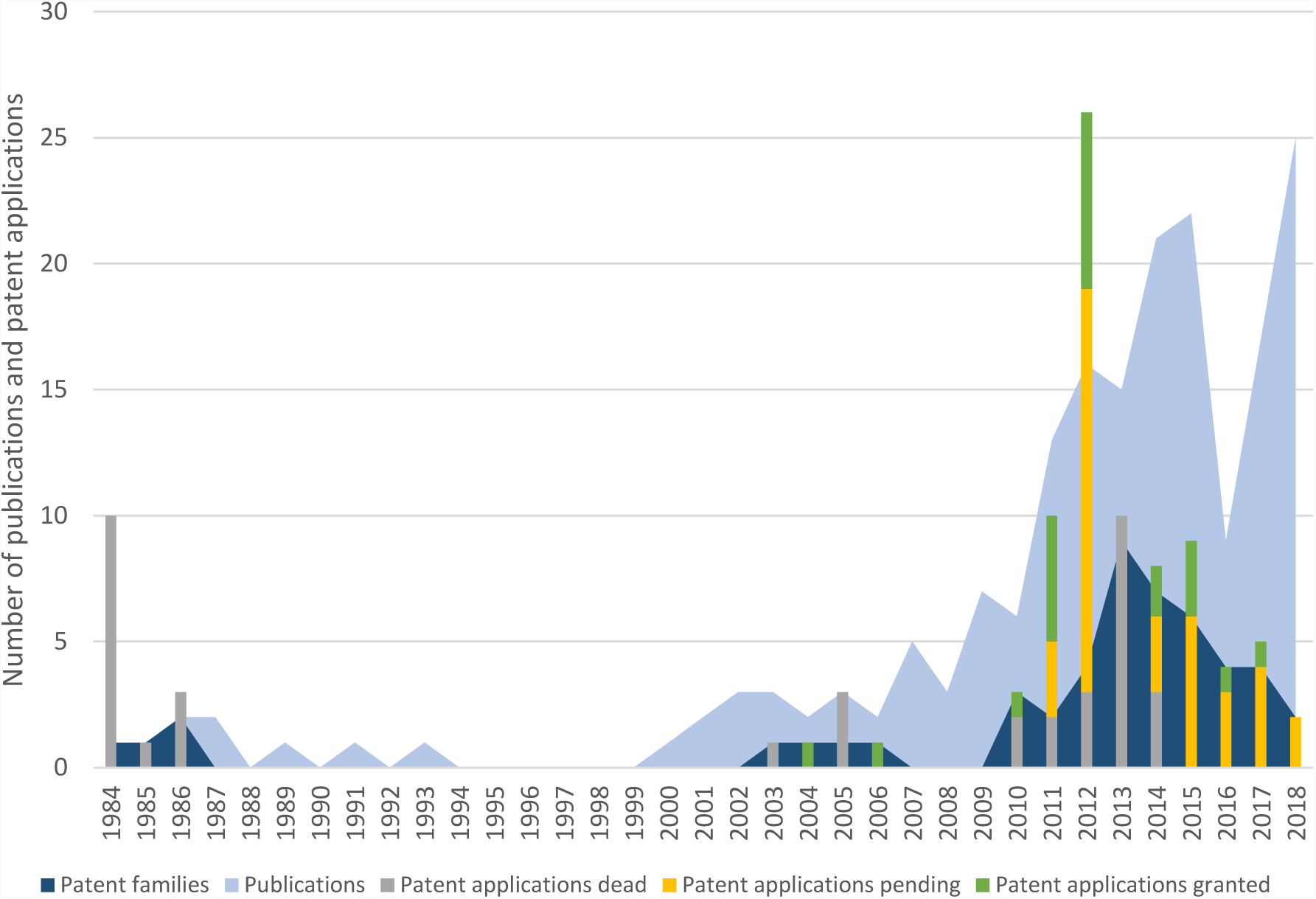
An overview of the number of scientific publications, patent families, patents and patent applications in the field of phage biocontrol in crop production organized by year. The light blue area chart represents the number of scientific publications and the dark blue area chart the number of patent families per priority year. The bars represent the number of patent applications per priority year: green (“Granted patents”) corresponds to the number of granted patents, yellow (“Pending applications”) pending applications and grey (“Dead applications and patents”) dead patents and applications. This last group consists of patents and applications that are abandoned, refused, withdrawn, deemed to be withdrawn or patents that are lapsed.

In sharp contrast, 137 scientific publications (1984-2019) were found within Web of Science and PubMed (Figure 1 – light blue area chart). Publications on phage biocontrol were scarce in the eighties and nineties of the 20^th^ century. However, from the year 2000 onwards, this number has increased steadily (25 peer reviewed publications in 2018). In other words, there is an increasing trend in the number of publications over the past decade, demonstrating a discrepancy between scientific publishing and patent filing. However, not all patent applications from 2017, 2018 and 2019 are publically available as the 18 month period before publishing is not passed yet.

When looking at the patent documents in more detail (Figure 1 – bar charts), a distinction should be made between “Granted”, “Pending” and “Dead”, based on their legal status as derived from Espacenet. In total, 59 patents and patent applications (61%) are active – 22 patents (23%) have been granted, 37 applications (38%) are pending – and 38 patent documents (39%) can be considered dead. From the latter 39% are applications that were deemed to be withdrawn, 16% are rejected applications and 16% are lapsed patents. Figure 1 shows an overview of the percentages granted, pending and dead documents with the same priority year. The first active patent from the dataset dates from 2004 (JP4532959 (B2)). In 2011 there were two families filed, represented by GB20110010647 and JP20110102153, containing in total six patents and four applications respectively. All the applications and patents within the first family remain active – 50% is granted and 50% is pending – whereas two out of four patent applications from the latter family have been rejected. 2012 has the most patents, 26 in total, divided among four different families represented by GB20120017097, KR101584214, MX2012011440 and US201261716245. This last family contains four granted patents and sixteen applications. The family represented by GB20120017097 consists of one pending and two dead applications and one granted patent (US9278141 (B2)). In 2013, there was a peak in the amount of patent families filed as the number reached nine families. These families consist in total of ten applications which are all dead. The majority of these applications were deemed to be withdrawn (80%), meaning that the designation fee was not paid.

### Applicants and geographical distribution

Analyzing the applicants of the different patent documents, it shows that 56% (54) of the 97 patent documents have been filed by academia, whereas 37% (36) are linked to industry (without joint applicants) and 7% (7) are joint applicants. This means that the family, or patent documents belonging to a family, have more than one applicant and thus the rights of the patent are distributed among the different partners. The most prominent academic applicants based on the amount of patent families filed by these applicants are the University of Hiroshima (JP) (21% of patent families - 6/28) and the Rural Development Administration (KR) (14% - 4/28). The most prolific company in terms of patent filing is Qingdao Biological Technology Corporation LTD (CN), accounting for 50% (8/16). However, it is worthwhile to mention that all patents within the database from the Qingdao Biological Technology Corporation LTD have been withdrawn. Remarkably, only 4 out of 21 granted patents are filed by applicants from the private sector and all of these patents belong to the same family (represented by GB20110010647 – Fixed Phage).

Figure 2 provides an overview of the relative contributions by countries in terms of filed and granted patent applications. 1% of all patent documents were filed in Africa (ZA), 3% in Oceania (AU – NZ), 13% in Europe (DE – EA – EP – ES – GB – HU – IT) and 13% worldwide. The majority of the patent documents have been filed in North America and Asia: 16% in the USA and Canada and 43% in Asia. These Asian documents consist of 45% (19/42) Chinese, 31% (13/42) Japanese, 21% (9/42) Korean and 2% (1/42) Indian applications and patents. When analyzing the legal status of the different patents organized per continent, 33% (14/42) of the Asian patents have been granted, 26% (11/42) of the applications are pending and 40% (17/42) is dead. The large portion of dead patents and applications is also visible among the European – 54% (7/13) – and among the North American documents – 44% (7/16). Only 8% (1/13) of European patents are granted, while this is 25% (4/16) of the North American patents. The amount of pending applications is similar as 38% (5/13) and 31% (5/16) of the European and North American applications are pending, respectively. On the other hand, in case of the South American patent applications, 89% (8/9) is pending and 11% is dead. Notably is that 7/9 of these South American patents are part of the same patent family (US2016309723).

**Figure 2.**
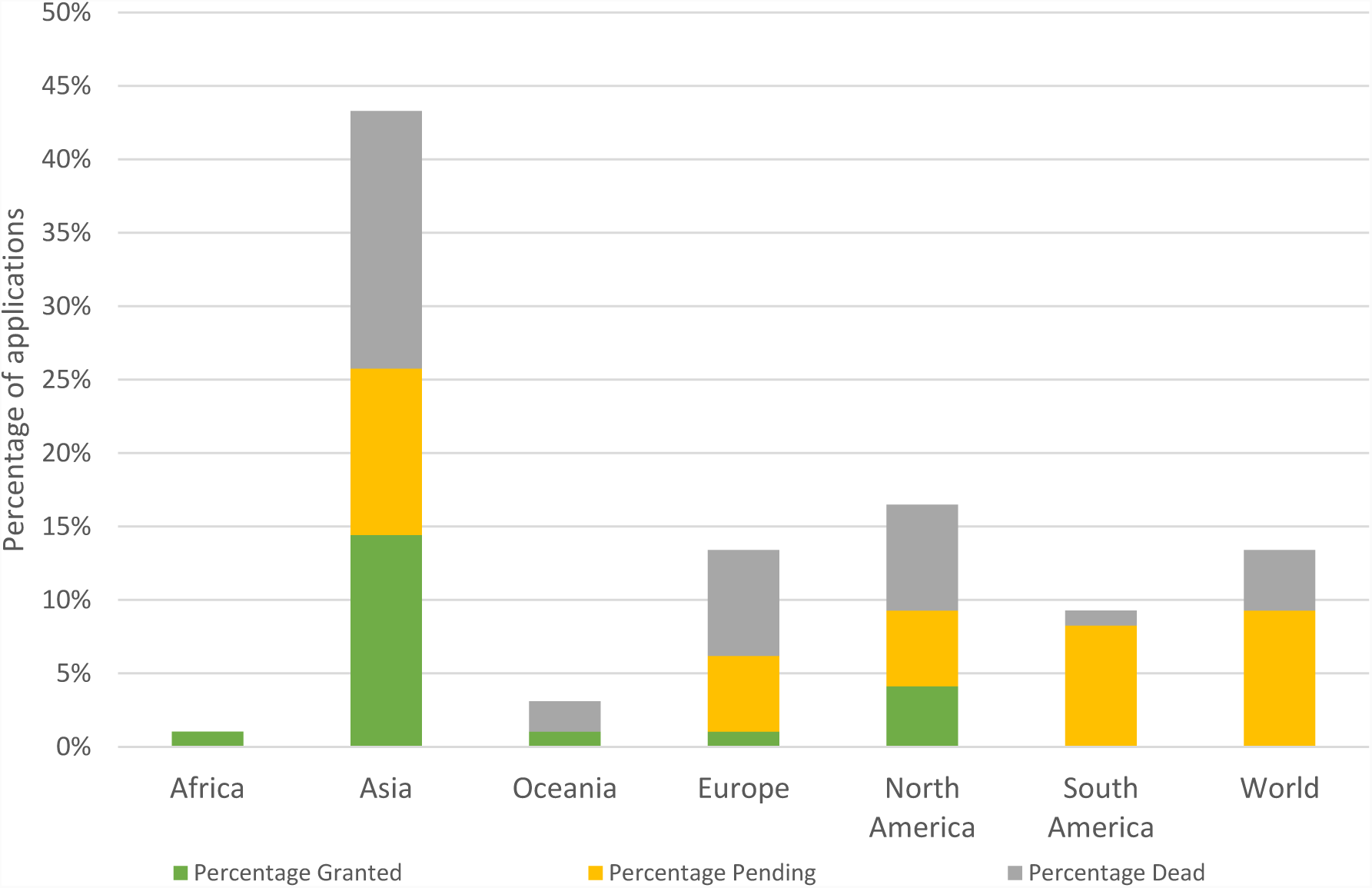
Percentage of patents and applications organized per continent. The total height of the bars indicate the percentage of patents and applications per continent. Africa (ZA), Asia (CN, IN, JP, KR), Oceania (AU, NZ), Europe (DE, EA, EP, ES, GB, IT), North America (CA, US), South America (AR, BR, CL, CR, GT, MX, PE) and world applications. In green the percentage of granted patents, in yellow the percentage of pending applications and in grey the percentage of dead applications and dead patents.

Furthermore, China has the highest percentage of dead patents with 63% (12/19). The majority of these patent were deemed to be withdrawn after the admission fee was not paid (92%). On the contrary, Japan and Korea have the highest amount of granted patents, 38% (5/13) and 67% (6/9) of the patents filed are granted and still active respectively.

### Categorization of patents and scientific publications

As the most prominent bacterial species belong to the genera of *Agrobacterium, Dickeya, Erwinia, Pectobacterium, Pseudomonas, Ralstonia, Xanthomonas* and *Xylella* [9], patent families and scientific publications have been categorized and quantified among these groups according to (in)dependent claims (patent applications) and abstracts (scientific publications) (Figure 3). Combinations of genera have been created when phages against certain pathogens are combined in a single cocktail (*Dickeya / Pectobacterium* and *Xanthomonas / Xylella*). A patent family or publication was classified as “Other” if it concerns a phage infecting bacteria of other genera (e.g. *Streptomyces* – “McKenna, F., et al. 2001. Novel In vivo use of a polyvalent Streptomyces phage to disinfest Streptomyces scabies-infected seed potatoes. Plant Pathol. 50:666-675.*”*) or if the paper/publication has no specification of the type of phage nor in the independent nor in the dependent claims (e.g. WO2016154602 – “A method of preparing a phage composition for killing or degrading fitness of a pest”).

**Figure 3.**
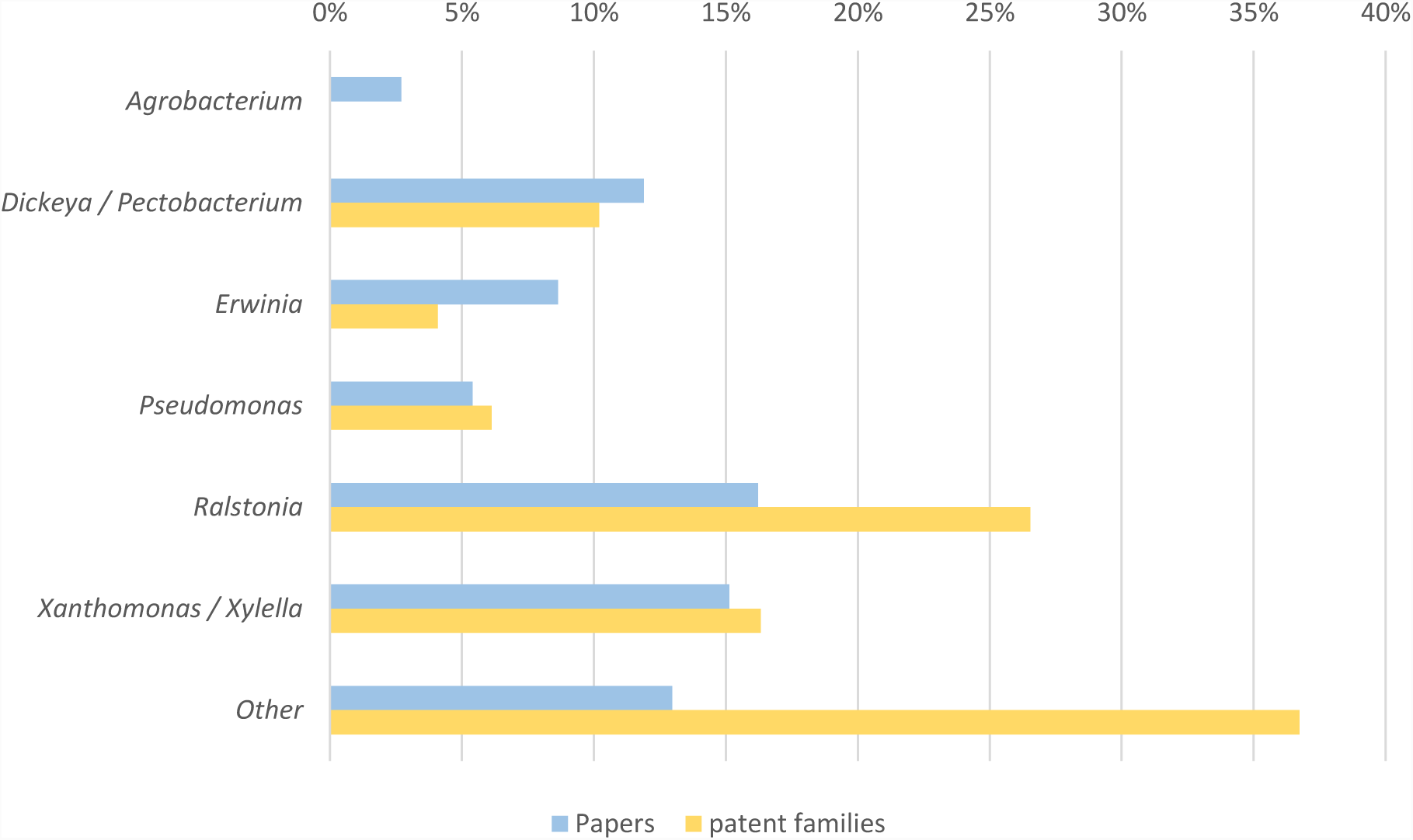
Distribution of patent families and scientific publications classified according to the bacterial genera that is tackled by the phage product. Based on Mansfield *et al.,* 2012 seven categories (*Agrobacterium, Dickeya / Pectobacterium, Erwinia, Pseudomonas, Ralstonia, Xanthomonas / Xylella* and Other) have been made to classify patent families and publications. Groupings of bacterial genera (*Dickeya – Pectobacterium* and *Xanthomonas – Xylella*) were created as phage cocktails are created to tackle both bacterial genera. The last category “Other” consists of patent families and publications that do not specify the bacterial pathogen that is being targeted or that tackle an alternative bacterial genera.

The least researched bacterial genera within both databases are *Agrobacterium* (0% patent families, 3% publications), *Pseudomonas* (6% patent families, 5% publications) and *Erwinia* (4% patent families, 9% publications). The most represented bacterial genera within the datasets as suggested by Figure 3 are *Ralstonia, Xanthomonas* - *Xylella* and *Dickeya* - *Pectobacterium*. 16% of the scientific publications discuss phages infecting *Ralstonia*, 15% *Xanthomonas, Xylella* or a combination of *Xanthomonas and Xylella* and 12% *Dickeya, Pectobacterium* or a combination of *Dickeya* and *Pectobacterium.* In case of the patent applications, a similar trend can be observed: 27% *Ralstonia*, 16% *Xanthomonas* / *Xylella* and 10% *Dickeya* / *Pectobacterium*.

A closer inspection of the different categories reveals that three patent families within the category of *Ralstonia* have included a private company as co-applicant (23% - 3/13). On the other hand, inventions from all patent families filed by public institutes have also been published in scientific papers (Supplementary Table 3). In case of the category of *Xanthomonas* / *Xylella*, a similar trend can be observed, as there are three families filed by industrial applicants represented by WO2015200519 by Auxergen (US), US2017142976 by Fairhaven Vineyards (US) and HU1700178 Enviroinvest Koernyezetvedelmi es biotechnologiai (HU).

The other families are filed by the University of Hiroshima (JP), Texas A&M University (US), University Huazhong (CN) and the National institute for Agro-Environmental Sciences (JP), all situated in the public sector. This is also the case for *Dickeya* and *Pectobacterium* as no patents were filed by private institutes. The applicants here include the Rural Development Administration (KR) and the national university of Seoul (KR). The majority of the families filed by private institutes are classified as “Other” (37%), indicating that the patents are not discussing the major genera of bacteria but rather do not specify the phages nor their host. Hence the claims are defined broadly. Within this group, the majority of the families are filed by industrial applicants (88%): Fixed phage (GB), Epibiome (US), Qingdao Biological Technology Corporation LTD (CN) and Internalle (US).

### Claim and abstract analysis

A claim analysis has been performed to determine the scope of the individual patents along with an analysis of the scientific publications. Among the dataset of 91 patent documents analyzed, 21 were active, granted patents. In total, 79 independent claims were systematically analyzed and classified in five different categories (Figure 4): (1) Phage (e.g. *“A bacteriophage able to lyse cells of Ralstonia solanacearum selected from the group of the following: a) vRsoP-WF2 (DSM 32039), vRsoP-WM2 (DSM32040), vRsoP-WR2 (DSM32041), or b) a podovirus whose genome has the sequence of SEQ ID NO: 1 (corresponding to vRsoP-WF2), SEQ ID NO: 2 (corresponding to vRsoP-WM2) or SEQ ID NO: 3 (corresponding to vRsoP-WR2)”* - ES2592352), (2) Composition (e.g. “*A composition for inhibiting or preventing the growth of Pectobacterium carotovorum, which comprises as an active ingredient bacteriophage PM-2 (KACC97022P) having an entire genome sequence consisting of the nucleotide sequence of SEQ ID NO: 1.”* – KR101797463), (3) Production (e.g. “*A method of propagating a virulent bacteriophage that includes X fastidiosa in its host range, comprising the steps of: (a) infecting a culture of Xanthomonas bacteria with said virulent bacteriophage; (b) allowing said bacteriophage to propagate; and (c) isolating virulent bacteriophage particles from the culture.” – “*Production claim” - AU2013331060) and (4) Treatment (e.g. “*A method of preventing or reducing symptoms or disease associated with Xylella fastidiosa or Xanthomonas in a plant, comprising contacting said plant with particles of at least one virulent bacteriophage, wherein Xylella fastidiosa and/or Xanthomonas axonopodis are hosts of the bacteriophage, wherein the bacteriophage is a Xfas 300-type bacteriophage and displays the following characteristics: (a) the bacteriophage is capable of lysing said Xylella fastidiosa and/or Xanthomonas bacteria; (b) the bacteriophage infects a cell by binding to a Type IV pilus; (c) the bacteriophage comprises a non-contractile tail with a capsid size ranging from 58-68 nm in diameter and belongs to the Podoviridae family; (d) the genomic size of the bacteriophage is about 43300 bp to 44600 bp; and (e) the bacteriophage prevents or reduces symptoms associated with Pierce’s disease in a plant or plants.”* – US9357785). The same categories were used to classify the scientific publications based on the abstract of the publication. Both scientific papers and patents can fall in different categories as it may discuss one of more phages and their basic characterization, the composition of a cocktail and the testing of this cocktail in bioassays and/or field trials. As Figure 4 shows, there are differences between the relative contributions of scientific papers and patents that address a specific topic. The majority of the scientific papers focus on the isolation and basic characterization of one or more phages (84%). 36% of the manuscripts discuss the composition of a phage cocktail and 39% use this cocktail in bioassays and/or field trials. The least represented topics the production methods of phage (4%). In contrast, the majority of the patents (90%) protects the use of phages to treat a plant in one way or another. 76% of the patents claim the composition of a product containing phage, 62% protect the phage itself and 24% contains claims protecting the production of the phage product. On the other hand, no patents within the database contain claims to protect methods of detection nor of the phage nor of the host.

**Figure 4.**
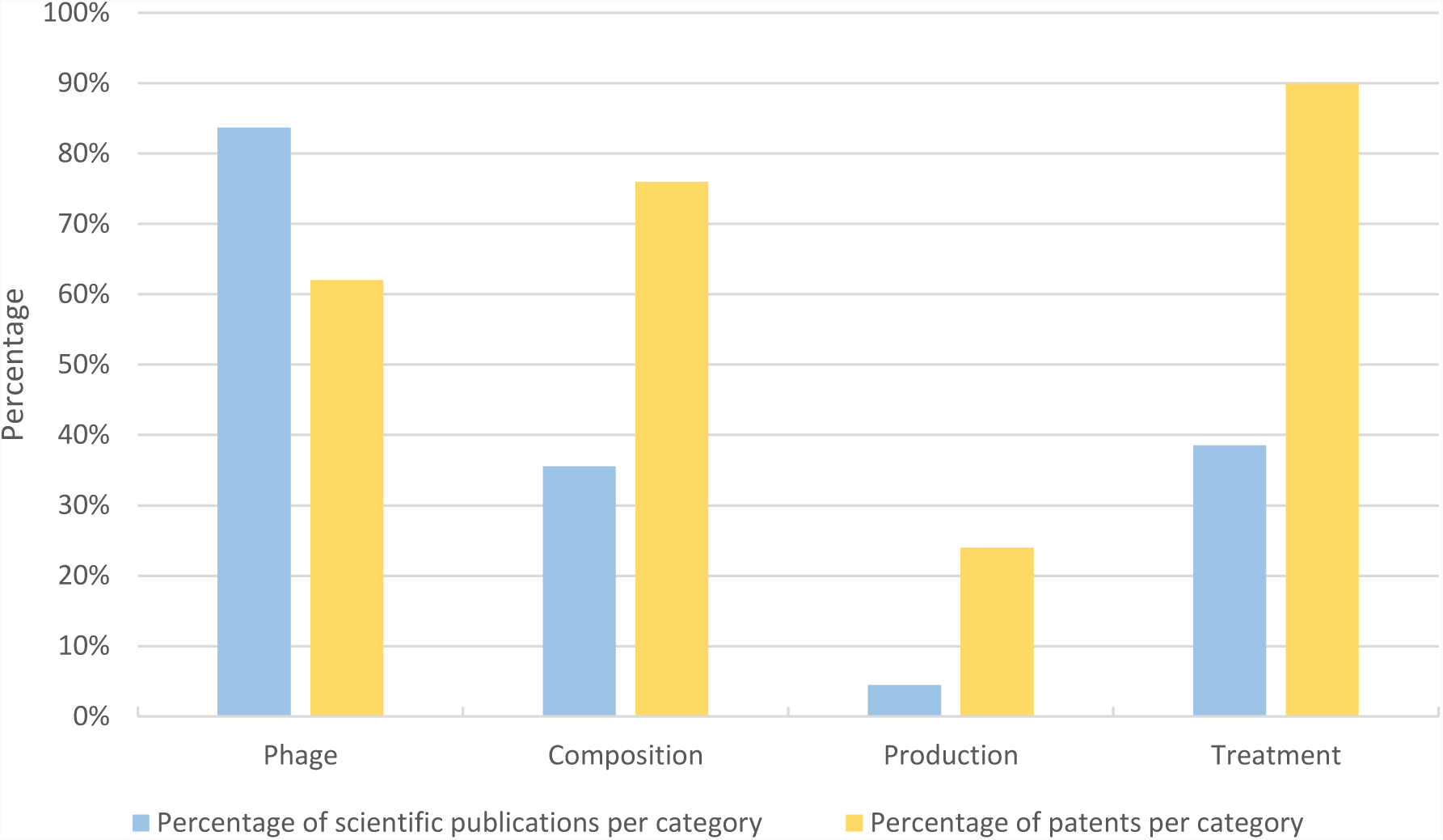
Claim and abstract analysis of the active, granted patents and scientific publications. In total, 79 independent claims from 21 patents and 137 abstracts from scientific papers were categorized among five different categories: (1) Phage – here the phage was described as the active ingredient or the isolation of a phage was described, (2) Cocktail – this category contains claims that protect the combination of phages and publications that describe a phage cocktail, (3) Production – ways of how the phage is produced and (4) Treatment – claims that protect the use of phages to fight a specific bacterial infection or methods and application strategies for using the phage (e.g. bioassays, field trials). In blue the percentages of publications are shown, in yellow the percentage of claims. Note: one publication and patent can be categorized in multiple categories.

When evaluating the 77 independent claims of all active, granted patents, the majority of the claims are process claims protecting different methods to use phages as a treatment for bacterial infection (40% - Supplementary Figure 1). Furthermore, this data shows that 30% of the claims protect the composition of the phage product. This means that a combination of phages is protected and/or the formulation of the final product. 19% can be considered as compound claims as the claim is protecting the phage(s) as an active ingredient and only 10% of the claims are production claims.

The 77 independent claims have also been categorized within three classes: narrow, intermediate and broadly defined. This gives an indication whether a specific claim can easily be circumvented (narrowly defined) or not (broadly defined). Table 1 gives an overview of all the granted patents and the independent claims that belong to this patent. 26% of the claims are narrow claims (e.g. “*The invention relates to a method for applying a bacteriophage of R. solanacearum, characterized in that the bacteriophage liquid of R. solanacearum is placed in a sterile syringe needle obliquely inserted into the stem of the tobacco plant, and then the bacterial phage liquid is covered with sterile mineral oil to prevent evaporation. And pollution, so that the R. solanaceans phage directly enters the stem of the tobacco through a sterile syringe needle; the sterile mineral oil is prepared by pouring 300 ml of mineral oil into a 500 ml screw bottle and sterilizing at 121 ° C 30min, then stored and reserved after cooling; the sterile syringe needle is prepared by using a sterile blister needle overnight, rinsing twice with sterile water, drying at 50 ° C after autoclaving, and after cooling to room temperature, Packed and stored for use; each tobacco stem is injected with 50-100 μl of R. solanacearum phage solution; in the sterile syringe needle inserted into the stem of the tobacco plant, the amount of sterile mineral oil is 50-100 μl” – “*Treatment claim” - CN104542717). 30% of the claims are intermediate as they are do not cover specific details on the invention (e.g. “*A method for controlling bacterial wilt disease bacteria, which comprises spraying the bacteriophage strain according to any one of claims 1 to 3 to a plant or soil.”* – “Treatment claim” - JP4532959). Finally, 44% can be defined as broad claims (e.g. “*An isolated bacteriophage which is toxic to X. fastidiosa and Xanthomonas species, wherein said bacteriophage is at least one member selected from the group consisting of Xfas 100 phage type bacteriophage and Xfas 300 phage type bacteriophage Of the bacteriophage, wherein the Xfas 100 phage type is selected from the group consisting of SEQ ID NO: 11, SEQ ID NO: 12, SEQ ID NO: 13, SEQ ID NO: 14, SEQ ID NO: 15, SEQ ID NO: 16, SEQ ID NO: 17, and SEQ ID NO: 18 Wherein the Xfas 300 phage type comprises a genome having a DNA sequence that is 99% or more identical to the sequence of SEQ ID NO: 19, SEQ ID NO: 20, SEQ ID NO: 21, SEQ ID NO: 22, SEQ ID NO: 23 and SEQ ID NO: 24 Select from It comprises a sequence a genome having the same DNA sequence 99% or more that is, the bacteriophage.” – “*Phage claim”-JP6391579).

**Table 1.** Claim analysis of the granted patents. Within this table, all granted patents have been organized according to the bacterial genera they describe, their patent number, the applicant (private sector blue and public sector yellow) and the nationality of the applicant (CN China, ES Spain, GB United Kingdom, JP Japan, KR Korea and US United States) and the filing year. Moreover, the claims have been categorized according the four different categories: (1) Phage: claims protecting one or multiple phages and their genome sequence, (2) Composition: claims discussing the composition of a phage product, (3) Production: methods to produce phages and (4) Treatment: methods to use phages as a treatment for plant diseases. If a patent contains claims that belong to a specific category, it is quantified. The colors indicate the scope of a claim or set of claims: very narrow, intermediate or broadly defined (green, orange and red, respectively).

| BACTERIAL<br>GENERA | PATENT<br>NUMBER | APPLICANT | FILING<br>YEAR | TYPE OF CLAIM |  |  |  |
| --- | --- | --- | --- | --- | --- | --- | --- |
|  |  |  |  | Phage | Composition | Production | Treatment |
| OTHER | US9539343 | Fixed Phage (GB) | 2012 |  | 1 |  | 1 |
|  | CN103747792 | Fixed Phage (GB) | 2012 |  | 1 |  | 1 |
|  | JP6230994 | Fixed Phage (GB) | 2012 |  | 1 |  | 1 |
|  | US9278141 | Fixed Phage (GB) | 2013 |  | 5 |  |  |
|  | CN104630154 | University of Zhejiang (CN) | 2015 | 1 | 1 | 1 | 2 |
|  | KR101887987 | Pukyong National University (KR) | 2016 | 1 | 1 |  | 1 |
| DICKEYA/<br>PECTOBACTERIUM | KR101368328 | Rural development Administration (KR) | 2010 | 1 | 1 |  |  |
|  | KR101790019 | Rural development administration (KR) | 2014 | 1 | 1 |  | 1 |
|  | KR101797463 | Rural development Administration (KR) | 2014 | 1 | 1 |  | 1 |
|  | KR101891298 | Seoul National University (KR) | 2016 | 1 | 2 |  | 1 |
| **PSEUDOMONAS** | KR101584214 | Chungbuk National University (KR) | 2012 |  | 1 |  | 1 |
| **RALSTONIA** | JP4532959 | Sanin Kensetsu Kogyo (JP) | 2004 | 3 | 2 |  | 2 |
|  | JP4862154 | University of Hiroshima (JP) | 2006 | 1 | 1 |  | 3 |
|  | US9380786 | University of Hiroshima (JP) | 2012 |  |  |  | 1 |
|  | JP5812466 | University of Hiroshima (JP) | 2011 | 1 |  |  | 3 |
|  | CN104542717 | Guizhou Tobacco science research institute (CN) | 2015 |  |  | 1 | 1 |
|  | ES2592352 | University of Valencia (ES) | 2015 | 1 | 1 |  | 2 |
| **XANTHOMONAS/**<br>**XYLELLA** | AU2013331060 | Texas A&M University (US) | 2013 | 1 | 1 | 2 | 4 |
|  | JP6391579 | Texas A&M University (US) | 2013 | 1 | 1 | 2 | 2 |
|  | US9357785 | Texas A&M University (US) | 2013 |  |  |  | 1 |
|  | ZA201502230 | Texas A&M University (US) | 2013 | 1 | 1 | 2 | 2 |

The claims categorized as “Phage claims” are generally narrow defined (41%) as the claims protect the phage and its genomic sequence. However, highly similar phages with differences in their genomic sequence fall outside the scope of these claim. Thus, the claims can be circumvented. A judge, however, may interpret by the claim as equivalent by doctrine of equivalence. The isolation of a different phage that targets the same bacterium also circumvents the claim. On the other hand, the majority of the “Composition claims” and “Production claims” are difficult to circumvent as they are broadly defined (56% and 75% respectively).

As mentioned above, four out of 21 granted patents have been filed by a private organization (Fixed Phage) and these patents belong to the same patent family. When looking at the claims for these patents, one notices that all the claims are broadly defined. As the invention discusses the use of phages in a composition to fight bacterial infections without defining the phage itself, the majority of the claims protect the composition of the product rather than the phage or the production of the product.

## 4. Discussion

Within this study, we analyzed publishing and patenting activities in phage biocontrol as a crop protectant by using the number of scientific papers and patent documents as a measuring tool for assessing interest in this area. Furthermore, we looked into the applicants, geographical distribution, legal status and scope of the patent documents by categorizing both the patents and scientific manuscripts by claims and abstracts. This allowed us to make a general comparison between the focus of patents and scientific literature and the willingness of R&D to look into the potential of phages as a part of IPM.

### Despite a growing interest in phage biocontrol, patenting activities remain limited

The increasing number of scientific publications (Figure 1) shows that there is indeed a growing interest to use phages as an alternative to existing plant protecting products. The number of patent filings seems to fall behind in 2016 but increases slightly in 2017. However, it is premature to determine a trend since not all the applications filed in 2017 and 2018 have passed the 18 months period before publication and thus are not made publicly available [24]. 56% of the patent documents were filed by non-profit organizations like the University of Hiroshima (JP), The Rural Development Administration of South-Korea (KR) and Texas A&M University (US). This confirms the slightly stronger interest from non-profit organizations to protect phage biocontrol inventions compared to industry. It might also suggest that private companies are still dangling in the start-up phase or have not picked-up the topic yet (apart from a few early adopters). This could also indicate that the expertise still remains in non-profit and still needs to be transferred to the private sector.

The assessment of the legal status of the documents shows a high fraction of dead documents (39%). Within this group of dead applications, 16% of the applications have been rejected as the invention was not considered novel (US2009053179) and/or did not include an inventive step (JP2005073562). The majority of the dead documents are applications that were deemed to be withdrawn (39%) and patents that are lapsed (16% - no oppositions were filed e.g. EP0182106). In both cases, the applicants did not pay the fees needed to maintain the rights of the patent. For some patent applications this is due to a negative search report (GB2519913 and WO2007044428) in which the invention was not found novel nor inventive. On the other hand, not paying the fees may also imply internal shifts of interest for commercial or other reasons. Hurdles in the regulation of biopesticides may influence such decisions as well. In Europe, for instance, the registration exists of two phases: registration of the active ingredient at European level by the European Food Safety Authority (EFSA) and next the authorization of the formulated product at member state level leading to bureaucratic difficulties [17]. As phages are highly specific and can only infect a few strains of one bacterial species, they are often combined in a cocktail [25]. In terms of registrations, this means that every phage in the cocktail should be registered as an active ingredient (Regulation (EC) No 1107/2009) and reformulations of the product require reregistrations [4]. The cost of such registrations may be high (e.g. the registration costs of products corresponding to a “New Active Ingredient, Non-food use; outdoor; reduced risk” can go up to $436,004) [26]. Moreover, phage genomes are variable due to evolutionary fluxes [27], thus phages cannot be considered as stable, fixed products which could impede registration [28]. Luckily, this can partially be addressed by recent insights on genome-based taxonomy that phages sharing 95% nucleotide similarity are considered as isolates the same species [29]. Changes in the regulation including fast track registration, priority registration and zonal authorization (*i.e.* authorization for all of the EU instead of registration per country) could function as an incentive to further develop innovations and promote patenting in biocontrol [30]. These reasons could also explain the low number of filed applications in Europe and North America as observed in Figure 2. Nevertheless, in the USA, a first phage product line, AgriPhage™, was registered in the US (2005) by Omnilytics (part of Phagelux). AgriPhage™ is approved by the Environmental Protection Agency (EPA) and contains products with phages against *Xanthomonas campestris* pv. *vesicatoria* and *Pseudomonas syringae* pv. *tomato.* Currently, Omnilytics has added cocktails against *Erwinia amylovora* and *Clavibacter michiganensis* subsp. *michiganensis* to their product line [31]. Notably, none of these products or phages have been found by the authors in the public assessed databases. This may imply that the patents could be licensed (e.g. from an academic partner), the applications are not publicly available yet or the product is not patented opening the discussion whether patenting is indeed crucial for commercial activities of phage applications in agriculture.

### Efforts in Asia to protect phage biocontrol preparations

The geographical distribution of the patent documents (Figure 2) shows that the majority was filed in Asia (43%). This observation might be explained by local governmental stimuli to promote patenting to cause rapid economic growth [32,33]. China, for instance has implemented so-called patent promotion policies. These policies stimulate patenting since tax incentives and subsidies are linked to patent ownership and have caused a booming growth of the number of Chinese patent applications and patents [33]. Furthermore, Asia in general, is taking measures to reduce the amount of chemical pesticides and fertilizers. China for instance has launched a “national research program on reduction in chemical pesticides and fertilizers in China” ($ 340 million) [30]. Within the study presented here, 19% of all documents have been filed in China. In contrast, 63% of these documents are dead from which 92% are applications that were deemed to be withdrawn as fees were not paid. This might implicate that the applicants lost their confidence in the invention. The number of granted, active patents is high in Japan and South-Korea, further illustrating Asian efforts to develop new biocontrol strategies. A study from the Food and Agriculture Organization from the United Nations (FAO) and the Asia and Pacific Plant Protection Commission (APPPC) has shown that both Japan and South-Korea are promoting integrated pest management strategies by setting pesticide reduction targets (KR) and appointing IPM expert groups (JP) [34].

### Scientific publications and patent documents show a correlation in terms of targeted pathogens

To get a more profound idea about the focus of the patent and scientific literature, both datasets have been classified according to the most prominent bacterial genera causing plant diseases [9] (Figure 3). In general, the percentages of scientific publications and patent families discussing a specific bacterial genus correspond quite well. Within the datasets here presented, the most studied bacterial genus is *Ralstonia.* Not surprisingly as these bacteria cause disease in a wide range of cash crops like tomato, potato, tobacco, eggplant and banana [35]. In the dataset, tomato is the most researched crop. The majority of the documents categorized within this group have been filed by non-profit organizations rather than industrial applicants. This suggests that the actual interest from industry is still rather limited compared to academia. The second most prominent group is the combination of *Xanthomonas* and *Xylella.* Both pathogens can have a significant impact in crop production and hence it is not remarkable that these bacteria are largely represented in phage biocontrol research [26, 27]. Furthermore, the majority of the applications have been filed by the public sector. The *Pseudomonas* genus, including of *Pseudomonas syringae* and its pathovars, are less represented within both databases. This is striking as *P. syringae* pathovars are considered as one of the most important PPB causing disease in different crops (tomato, bean, kiwi, leek and others) [38]. As Figure 3 shows, there is a discrepancy within the “*Other*” group. This category combines documents that talk about other bacterial genera or do not specify the bacterium that is being addressed. The majority of the patents and patent applications have been classified in this category since these do not specify a particular bacterial genus. From these patent documents, we observe that all the granted patents filed by industrial applicants have chosen this option (Table 1). This might suggest that industry defines claims containing bacteriophages as broad as possible to get patent exclusive rights to any phage product that falls under the protection of the patent. By applying this strategy, companies are taking risks in their patenting strategy because these broad claims are more susceptible in terms of novelty anticipation that may invalidate them [39].

### Granted patents include broad claims

The main focus of the scientific publications and the granted patents indicate a difference in emphasis (Figure 4). While the majority of the manuscripts discuss the isolation of one or more bacteriophages and their basic characterization, the majority of the patents claim a method of treating a bacterial infection by means of a phage product. Only 39% of scientific papers address the latter although this could be considered as the ultimate goal of phage biocontrol. This illustrates that first efforts are made in the field but that there are still opportunities for further innovations.

Limited efforts have been made towards patenting detection strategies for and by phages in the field of phage biocontrol in crop production. Since phages are highly specific to a specific bacterium and can locate their host in a complex matrix of bacteria they can easily be used as a detection tool [40]. However, this study suggests that little published evidence is available done in this area by both the scientific community and the industrial early adopters as there are no claims protecting possible methods of detection and only a small minority (8%) of scientific papers discuss this matter.

The claims that belong to the “Phage” category consist of claims that protect a natural occurring phage. According to Art. 3 of the Biotechnology Directive 98/44/EC (European Commission), natural phages can be patented because the phage is isolated from the environment [21]. The techniques to isolate phages are however similar to the ones used in 1920 which makes the patentability of natural phages fragile [20]. Nevertheless, claims for isolated phages can be of great value since patents covering such claims may give the right to prevent others from using that particular phage. On the contrary, as phages are abundantly present in the environment, the chances of finding a similar, yet different phage exists which may circumvent the claim. However, as the literature on phages applicable for phage biocontrol increases, anticipation of novelty as patentability requirement of a particular phage may become an issue. In the US, the requirement of novelty is more complicated, as the phage will be excluded for patentability if on the one side it is known or used within the US, or on the other side published or patented wherever in the world [21].

Figure 4 also shows that there is limited evidence on the optimization of the production of phage cocktails based on both scientific and patent literature. One could argue that the production of phages is phage dependent (specific bacterial strains, media, temperature) and thus keeping the production of phages secret could be a valid strategy to maintain a competitive edge [18].

On the other hand, many patents claim the combination of phages or phages as part of a formulation. This is illustrated by Supplementary Figure 1. Claiming a combination of phages can act as a buffer to reduce the risk of resistance development of the target bacterium. Phages are known to have different infection strategies and highly diverse genomes making the chances that a certain bacterium develops resistance (by altering receptors, CRISPRs, restriction enzymes) against two or even three phages in one cocktail are theoretically small [4,41,42]. The strength of these claims can be questioned as minor changes to the cocktail could already be sufficient to circumvent claims.

Table 1 depicts in which categories the individual patents can be classified. Here it is clear that the different patents combine different types of claims. Combined with the scope of the claims, this might support the previous statement that companies employ a “throw everything at the wall to see what sticks” patenting strategy because of the uncertainty and the ignorance how to achieve a strong protection for the invention [18].

### Phages and other viruses as part of an integrated pest management strategy

To safeguard public health and minimize impact on the environment, traditional pesticides used in crop production are currently being stigmatized. This triggers the research community to evaluate new, alternative methods to deal with different kinds of pests like fungi, bacteria and insects. The use of relatively high concentrations of natural predators to eradicate a certain pest, also known as augmentative biocontrol, is gaining in popularity [30]. In this regard, bacteriophages and other viruses, like insect viruses and mycoviruses, are ideal candidates as biocontrol agents because they are sustainable, specific and do not leave residues on the crop. Moreover, they fit into the framework of integrated pest management where the use of biocontrol agents is heavily promoted [30]. Viruses as biocontrol agents are, however, sometimes overlooked in IPM [15], which indicates that more efforts need to be taken to integrate these viral strategies. This review shows that there is indeed a basis within the scientific community to investigate the potential of phages to be used as biocontrol agents. In 2018, a Horizon 2020 consortium has been established to investigate viruses and their potential to serve as a possible solution against pests both as probiotic and viral treatment strategy (https://viroplant.eu/). Initiatives like these together with more in depth research are needed to provide fundamental insights to close the gap between academia and industry in this matter and to stimulate the industry to invest in phage biocontrol.

## Author Contributions

Conceptualization, DH, RL, IH and JW; methodology, DH and IH; writing—original draft preparation, DH; writing—review and editing, RL, IH and JW.

---

**Supplementary Figure 1.**
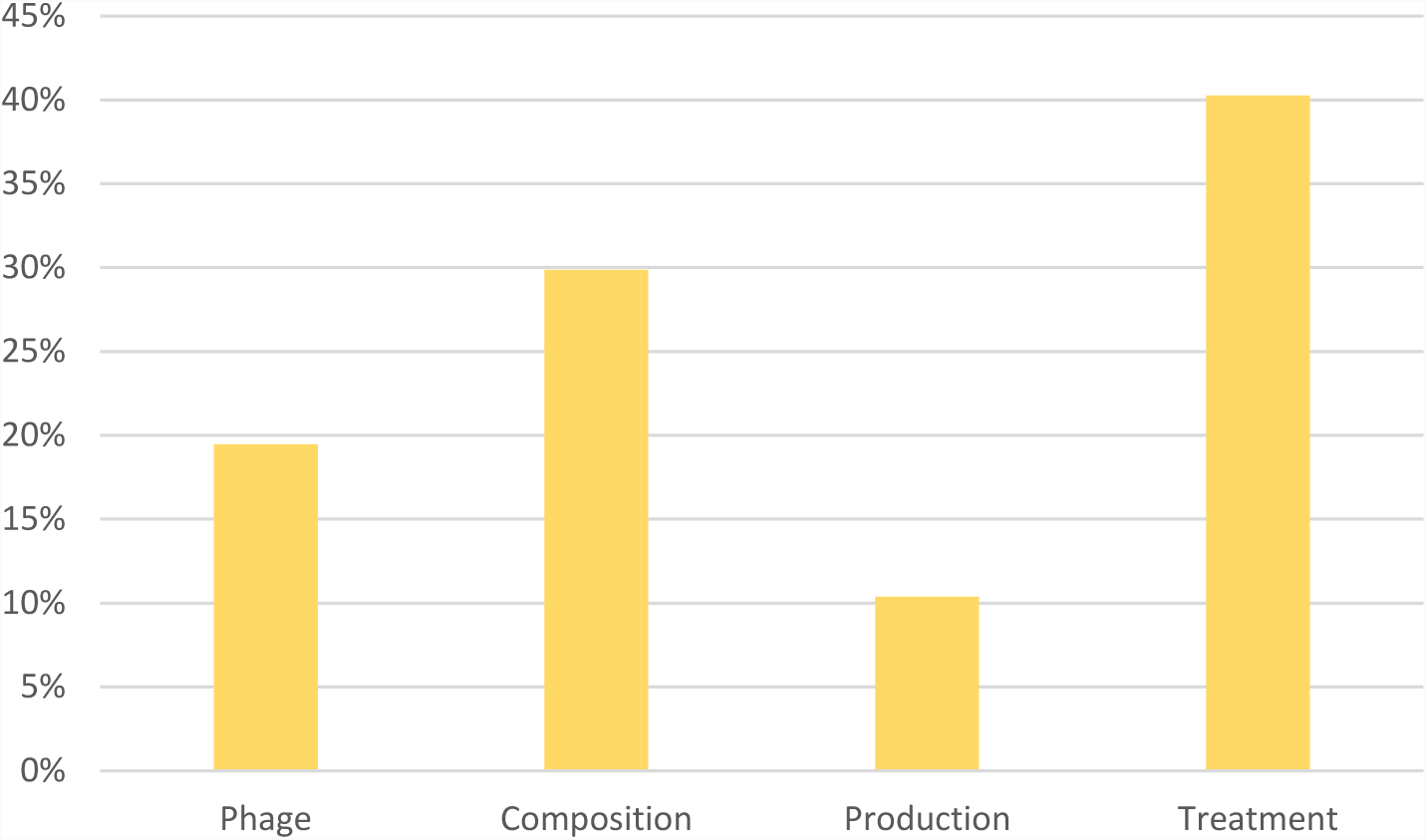
Percentages of independent claims per category. The independent claims, 77 in total, from the 21 granted patents have been categorized among five different categories: (1) Phage – here the phage was described as the active ingredient or the isolation of a phage was described, (2) Cocktail – this category contains claims that protect the combination of phages or phages as part of a composition (3) Detection – strategies to use phages for detection or to detect the phage itself, (4) Production – ways of how the phage is produced and (5) Treatment – claims that protect the use of phages to fight a specific bacterial infection or methods and application strategies for using phage.

## References

1. Twort, F.W. AN INVESTIGATION ON THE NATURE OF ULTRA-MICROSCOPIC VIRUSES. Lancet 1915, 186, 1241–1243.

2. D’Herelle, F.; Roux On an invisible microbe antagonistic toward dysenteric bacilli: brief note by Mr. F. D’Herelle, presented by Mr. Roux. Res. Microbiol. 158, 553–554.

3. Pirnay, J.P.; De Vos, D.; Verbeken, G.; Merabishvili, M.; Chanishvili, N.; Vaneechoutte, M.; Zizi, M.; Laire, G.; Lavigne, R.; Huys, I.; et al. The phage therapy paradigm: Prêt-à-porter or sur-mesure? Pharm. Res. 2011, 28, 934–937.

4. Buttimer, C.; McAuliffe, O.; Ross, R.P.; Hill, C.; O’Mahony, J.; Coffey, A. Bacteriophages and bacterial plant diseases. Front. Microbiol. 2017.

5. Mallmann, W., and Hemstreet, C. Isolation ofan inhibitory substance from plants. Agric. Res. 1924, 28, 599–602.

6. Okabe, N.; Goto, M. Bacteriophages of Plant Pathogens. Annu. Rev. Phytopathol. 1963, 1, 397–418.

7. Strange, R.; Scott, P.R. Plant disease: A threat to global food security. Annu. Rev. Phytopathol. 2005, 43, 83–116.

8. Inagro IWT - beheersing bacteriële pathogeen opkweek kolen-prei. Available online: http://leden.inagro.be/Wie-is-Inagro/Projecten/project/13960 (accessed on Dec 27, 2018).

9. Mansfield, J.; Genin, S.; Magori, S.; Citovsky, V.; Sriariyanum, M.; Ronald, P.; Dow, M.; Verdier, V.; Beer, S. V.; Machado, M.A.; et al. Top 10 plant pathogenic bacteria in molecular plant pathology. Mol. Plant Pathol. 2012, 13, 614–629.

10. Cooksey, D. a Genetics of Bactericide. Annu. Rev. Phytopathol. 1990, 28, 201–19.

11. Frampton, R.A.; Pitman, A.R.; Fineran, P.C. Advances in bacteriophage-mediated control of plant pathogens. Int. J. Microbiol. 2012, 2012.

12. Pietrzak, U.; McPhail, D.C. Copper accumulation, distribution and fractionation in vineyard soils of Victoria, Australia. Geoderma 2004, 122, 151–166.

13. Copping, L.G.; Duke, S.O. Natural products that have been used commercially as crop protection agents. Pest Manag. Sci. 2007, 63, 524–554.

14. Ray, D.K.; Mueller, N.D.; West, P.C.; Foley, J.A. Yield Trends Are Insufficient to Double Global Crop Production by 2050. PLoS One 2013, 8, e66428.

15. Stenberg, J.A. A Conceptual Framework for Integrated Pest Management. Trends Plant Sci. 2017, 22, 759–769.

16. McDougall Cost of Crop Protection Innovation Increases to $286 Million per Product Available online: https://www.ecpa.eu/news/cost-crop-protection-innovation-increases-286-million-product (accessed on Feb 9, 2019).

17. European Crop Protection Registering plant protection products in the EU; Brussels, 2013;

18. Todd, K. the Promising Viral Threat To Bacterial Resistance: the Uncertain Patentability of Phage Therapeutics and the Necessity of Alternative Incentives. J.D. Expect. 2019.

19. Williams, H.L. How do patents affect research investments? Annu Rev Econ. 2017, 9, 441–469.

20. Pirnay, J.-P.; Verbeken, G.; Rose, T.; Jennes, S.; Zizi, M.; Huys, I.; Lavigne, R.; Merabishvili, M.; Vaneechoutte, M.; Buckling, A.; et al. Introducing yesterday’s phage therapy in today’s medicine. Future Virol. 2012, 7, 379–390.

21. Verbeken, G. TOWARDS AN ADEQUATE REGULATORY FRAMEWORK FOR BACTERIOPHAGE THERAPY, KU Leuven and Royal Military Academy, 2015.

22. Verbeure, B.; Matthijs, G.; Van Overwalle, G. Analysing DNA patents in relation with diagnostic genetic testing. Eur. J. Hum. Genet. 2006, 14, 26–33.

23. Huys, I.; Berthels, N.; Matthijs, G.; Van Overwalle, G. Legal uncertainty in the area of genetic diagnostic testing. Nat. Biotechnol. 2009, 27, 903–909.

24. xEPO Basic definitions Available online: https://www.epo.org/searching-for-patents/helpful-resources/first-time-here/definitions.html (accessed on Feb 14, 2019).

25. Rombouts, S.; Volckaert, A.; Venneman, S.; Declercq, B.; Vandenheuvel, D.; Allonsius, C.N.; Van Malderghem, C.; Jang, H.B.; Briers, Y.; Noben, J.P.; et al. Characterization of novel bacteriophages for biocontrol of bacterial blight in leek caused by Pseudomonas syringae pv. porri. Front. Microbiol. 2016, 7, 1–15.

26. EPA PRIA Fee Category Table - Registration Division - New Active Ingredients Available online: https://www.epa.gov/pria-fees/pria-fee-category-table-registration-division-new-active-ingredients (accessed on Feb 15, 2019).

27. Barbosa, C.; Venail, P.; Holguin, A. V.; Vives, M.J. Co-Evolutionary Dynamics of the Bacteria Vibrio sp. CV1 and Phages V1G, V1P1, and V1P2: Implications for Phage Therapy. Microb. Ecol. 2013, 66, 897–905.

28. EuropeanCommission GUIDANCE DOCUMENT FOR THE ASSESSMENT OF THE EQUIVALENCE OF TECHNICAL GRADE ACTIVE INGREDIENTS FOR IDENTICAL MICROBIAL STRAINS OR ISOLATES APPROVED UNDER REGULATION (EC) No 1107 / 2009; 2014;

29. Adriaenssens, E.M.; Rodney Brister, J. How to name and classify your phage: An informal guide. Viruses 2017, 9, 1–9.

30. van Lenteren, J.C.; Bolckmans, K.; Köhl, J.; Ravensberg, W.J.; Urbaneja, A. Biological control using invertebrates and microorganisms: plenty of new opportunities. BioControl 2018, 63, 39–59.

31. AgriPhage TM Product info AgriPhage Available online: https://www.agriphage.com/product-info/ (accessed on Feb 9, 2019).

32. Erstling, J.; Strom, R. Korea’s Patent Policy and Its Impact on Economic Development: A Model for Emerging Countries. San Diego Int’l LJ 2009, 441–481.

33. Long, C.X.; Wang, J. China’s patent promotion policies and its quality implications. Sci. Public Policy 2018.

34. APPPC; FAO Plant protion profiles from Asia-Pacific countries - Chapter 3 Intergrated Pest Management Available online: http://www.fao.org/docrep/010/ag123e/AG123E22.htm (accessed on Feb 13, 2019).

35. Álvarez, B.; Biosca, E.G. Bacteriophage-Based Bacterial Wilt Biocontrol for an Environmentally Sustainable Agriculture. Front. Plant Sci. 2017, 8, 1–7.

36. Ryan, R.P.; Vorhölter, F.J.; Potnis, N.; Jones, J.B.; Van Sluys, M.A.; Bogdanove, A.J.; Dow, J.M. Pathogenomics of Xanthomonas: Understanding bacterium-plant interactions. Nat. Rev. Microbiol. 2011, 9, 344–355.

37. Janse, J.D.; Obradovic, A. XYLELLA FASTIDIOSA: ITS BIOLOGY, DIAGNOSIS, CONTROL AND RISKS. J. Plant Pathol. 2010, 92, S1.35–S1.48.

38. Xin, X.F.; Kvitko, B.; He, S.Y. Pseudomonas syringae: What it takes to be a pathogen. Nat. Rev. Microbiol. 2018, 16, 316–328.

39. WIPO IP and Business: Quality Patents: Claiming what Counts Available online: https://www.wipo.int/wipo_magazine/en/2006/01/article_0007.html (accessed on Feb 14, 2019).

40. Schofield, D.A.; Bull, C.T.; Rubio, I.; Wechter, W.P.; Westwater, C.; Molineux, I.J. Development of an engineered bioluminescent reporter phage for detection of bacterial blight of crucifers. Appl. Environ. Microbiol. 2012, 78, 3592–3598.

41. Labrie, S.J.; Samson, J.E.; Moineau, S. Bacteriophage resistance mechanisms. Nat. Rev. Microbiol. 2010, 8, 317–327.

42. Chan, B.K.; Abedon, S.T.; Loc-Carrillo, C. Phage cocktails and the future of phage therapy. Future Microbiol. 2013, 8.

